# Task demands shift motor learning from adaptation to feedback control in an unconstrained bimanual task

**DOI:** 10.1101/2025.05.01.651653

**Authors:** Rini Varghese, Cristina Rossi, Laura A Malone, Amy J Bastian

## Abstract

Bimanual actions are common in daily life, yet how task demands shape reliance on learning and control mechanisms remains largely unknown. We examined this in 73 participants who lifted a plate in virtual reality while we distorted visual motion of their right hand. During right-hand-only lifting, aftereffects were present but incomplete. Bimanual lifting with the same right-hand perturbation shifted learning further from adaptation toward feedback control: participants developed compensatory hand adjustments and showed smaller aftereffects. Relaxing precision demands improved success but neither restored aftereffects nor reduced compensation. Removing interlimb conflict by perturbing both hands did not improve success but eased compensatory adjustments and restored plate aftereffects to unimanual levels. Success rate during learning did not correlate with aftereffects, but faster movement speed did. This demonstrates that precision demands and interlimb conflict independently shift learning strategy in bimanual contexts.

## INTRODUCTION

The nervous system has many ways to flexibly control movement. Imagine a friend asks you to help move her prized encyclopedia collection. You agree, though you can’t imagine why anyone still keeps a full set. As you pick up the first box, it is heavier than expected. You don’t drop it, but the lift is slow and unstable; this is because you are correcting your movement based on feedback, which has inherent delays reaching the brain. For the next box you decide to use more force, getting it off the ground faster, though still not smoothly. A strategy like this is quick to deploy, but only a guess at the force required, not a precise estimate of the motor commands needed. As you move all the remaining boxes, it gets easier: you make fewer corrections and no longer think about how to lift, because the brain has learned to anticipate each box weight in advance, making the movement quick and accurate.

This example highlights three ways the nervous system adjusts motor commands when a movement does not go as planned: feedback-driven corrections, strategic compensation, and learning. Their balance depends on the context and constraints of the task. Here we ask to what extent adaptation—an error-driven form of learning—occurs during bimanual movements in a highly unconstrained setting.

Adaptation is driven by sensory prediction errors—the difference between the predicted and actual outcome of a movement.^1^ In our example, adaptation would operate if the boxes were predictable, all similar in size and weight. Errors are reduced across repeated lifts as the brain learns to predict the appropriate motor commands. In the laboratory, adaptation is studied by having people make repeated reaching movements against a perturbation, typically using a robot to push the hand off course (a force perturbation) or a display that makes an accurate movement appear off course (a visuomotor perturbation).^2,3^ Both induce sensory prediction errors, and people adapt their reaches gradually, trial by trial. A hallmark of adaptation is the aftereffect: when the perturbation is removed, people keep moving with the remapped pattern and must de-adapt over many trials, rather than simply switching it off.^4–6^

Less is known about adaptation during bimanual movements, even though most functional actions use both hands. The central question is how an error at one hand is distributed: corrections and adaptation can stay local or spread to the other hand. Planar bimanual experiments show this depends on task demands—when the hands jointly control one object, perturbing one hand can drive corrections ^7–13^ and adaptation in both,^14,15^ but when hands control separate objects, responses remain confined to the perturbed hand.^14^ This balance of shared versus independent control is graded, influenced by each hand’s contribution to the task^16, 17^ or the consistency of sensory predictions between hands.^18^ This flexibility means the motor system can adopt different strategies—adapting one hand predictively while the other relies on feedback corrections, or redistributing control asymmetrically across the hands—rather than being forced into a single solution.^19–21^

Yet despite such flexibility, how adaptation unfolds during less-constrained bimanual tasks remains largely unknown.^4^ Understanding how the nervous system balances adaptation versus other adjustments (e.g., feedback driven corrections and strategic compensations)^6,22^ may help design rehabilitation training methods, particularly if therapy is focused on compensatory mechanisms (e.g. strategy), better use of feedback control (e.g. sensory training), or learning new motor patterns.

We had participants play a virtual reality game in which they lift a virtual plate of grapes to feed a bird. Unbeknownst to the participant, we distorted visual feedback of the right-hand, so it appeared to move less than it really did. We first compared learning when the right-hand moved alone (unimanual) versus when both hands moved together (bimanual). We expected that adaptation would be incomplete given the unconstrained nature of the task. This would result in smaller aftereffects and a shift toward use of feedback control or strategic compensation. We predicted that there might be more adaptation in the unimanual case, because the bimanual case introduces interlimb conflict: when only the right-hand is perturbed, the left-hand supplies conflicting information about its true position, so participants may rely more on feedback control. We tested this in two ways. First, we relaxed the target precision requirements in the bimanual case, reasoning that larger aftereffects would emerge if participants no longer needed to rely as heavily on feedback control. Second, we removed the interlimb conflict by perturbing both hands. We hypothesized that wider targets or bilateral perturbation would each produce larger aftereffects, showing that each feature constrains how much adaptation the bimanual context expresses.

## RESULTS

We studied how task demands influence bimanual motor learning using a transport task in virtual reality that could be done unimanually or bimanually. Participants wore a Meta Quest 2 VR headset with real-time hand tracking via a LEAP Motion controller (Fig 1a) to lift a virtual plate of grapes to feed a bird on a target perch (Fig 1b). Upon donning the headset, the target position was automatically calibrated to each participant’s eye level based on the headset coordinates, which varied according to their seated height. The initial plate position was then set 30 cm directly below this individualized target position. Participants were told that the lift must move the plate into a visible yellow target zone at the end of the movement (Fig 1c). The basic experimental flow consisted of 260 trials across three phases with 30-second breaks after every 30 trials (Fig 1d): baseline phase (50 trials) with veridical visual feedback, learning phase (150 trials) where a visual gain applied to the y-position of the right wrist at plate contact was gradually reduced from 100% to 65% over 75 trials (ramp), then held constant for 75 trials (plateau), and washout phase (60 trials) where veridical visual feedback was restored to examine aftereffects.

**Figure 1.**
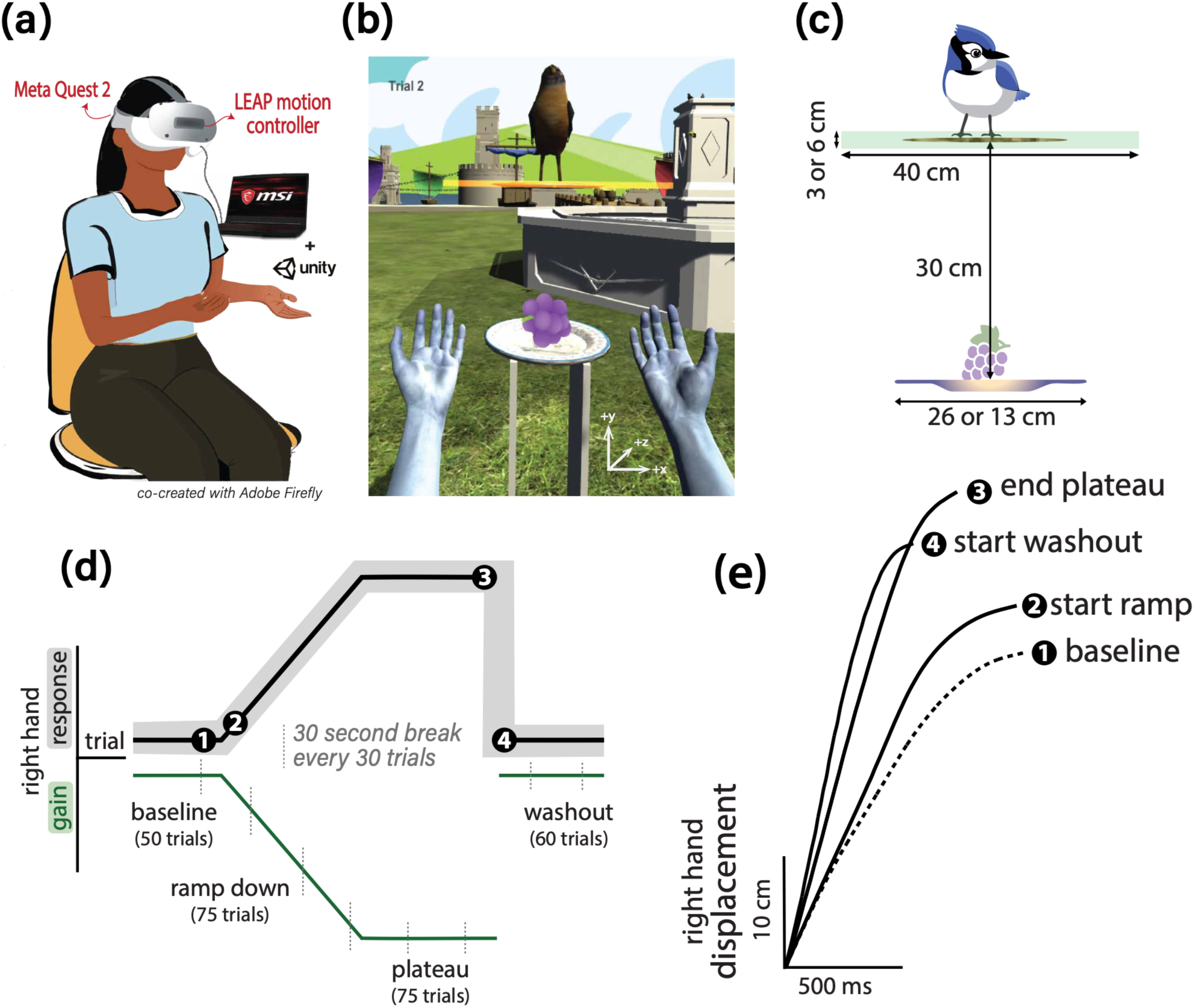
Setup and experimental paradigm. **(a)** Participants wore a Meta Quest 2 headset, on which a LEAP motion controller is mounted. **(b)** Game view shows real-time tracking and rendering of embodied hands and virtual objects. **(c)** Lifting task parameters including distance to target, target tolerance (narrow, 3 or wide, 6 cm in the two bimanual groups), and plate width (bimanual, 26 or unimanual, 13 cm). **(d)** Perturbation schedule showing gradual ramp down of visual gain of right-hand y-position to reach a maximum size of 65% of the ‘real’ right-hand’s y-position (green line). The dark gray line and shaded region are the expected responses and success zone in response to the gain perturbation. **(e)** Right-hand vertical displacement from one participant showing canonical response to the gain perturbation. Trace 1 is a baseline 30 cm lift. Trace 2 shows this subject lifting higher early in the ramp phase where the visual gain was beginning to change. Trace 3 shows the hand moving higher to counteract the full effect of the visual gain at the end of plateau. Trace 4 shows an example of the aftereffect in early washout (gain removed) where the hand moves higher than baseline, but not as high as trace 3. This smaller “aftereffect” compared to the end of plateau suggests that the participant adapted, but incompletely and may be using feedback control or strategy to correct the inflight movement.

Figure 1e shows timeseries plots of the upward right-hand movement (i.e., wrist) at different phases of the experiment. Trace 1 is a baseline 30 cm lift. Trace 2 shows this subject lifting higher early in the ramp phase where the visual gain was beginning to change. Trace 3 shows the hand moving higher and faster (steeper slope) to counteract the full effect of the visual gain at the end of plateau. Trace 4 shows an example of the aftereffect in early washout (gain removed) where the hand moves higher and faster than baseline, but not as high as trace 3. This smaller “aftereffect” compared to the end of plateau (trace 3) suggests that the participant adapted, but incompletely and may be using feedback control or strategy to correct the inflight movement.

For all experiments, we calculated s*uccess rates*. Success required correct *plate displacement (plate center in target zone)* for uni- and bimanual movements and correct plate alignment (right and left edges in target zone) for bimanual movements (see methods). Note that the plate size was smaller in the unimanual condition for more stable single hand control.

We then quantified the movement patterns that led to good performance. Participants could change their *right-hand displacement –* the vertical movement of the right-hand in response to the visual distortion during learning. They could also flex or pronate the right wrist slightly to raise the plate which we call *right tilt compensation*. In the bimanual condition, participants could additionally change their *left-hand displacement* and *left tilt compensation*.

Our primary adaptation metric was aftereffects in plate and right-hand displacement in early washout. This captured whether the movement pattern remapped to meet the task goal was retained when the perturbation was turned off and veridical feedback restored during washout. Smaller aftereffects in plate and right-hand displacement would suggest a shift toward feedback corrections. Aftereffects that are transient and rapidly return to baseline (e.g., 1-2 trials) rather than decaying gradually, would suggest a shift toward strategic learning that could be quickly abandoned.

Our original experiment compared a unimanual group (n = 20) to a bimanual group (n = 22), both performing lifts to a narrow (3 cm) target zone with a gain applied to the right hand. To examine what features of the bimanual task constrain adaptation, we recruited two additional groups that each isolated one component of the bimanual task. The bimanual-wide group (n = 16) experienced the same right-hand perturbation and target-alignment requirement as the bimanual group, but with a target zone twice the height (6 cm), giving more tolerance for alignment errors. The bimanual-both group (n = 15) kept the narrow (3 cm) target and alignment requirement but applied the gain perturbation to both hands rather than the right hand alone.

In the sections that follow, we first compare the unimanual and bimanual groups on performance, movement patterns, and adaptation (i.e., aftereffects), and then examine how widening the target or removing interlimb conflict altered these same measures.

### Success was limited by precision demands, not interlimb conflict

The unimanual group was more successful than the bimanual group at baseline, and success rates in both groups decreased as the perturbation ramped up (Fig 2a, triangles vs closed circles). By the end of learning, the unimanual group remained more successful than the bimanual group (mean difference = 38.5 ± 5.2%, z = 7.35, p<0.001; Fig 2b). Failures in all groups commonly occurred due to errors in plate displacement, alignment, or dropping the grapes or plate.

**Figure 2.**
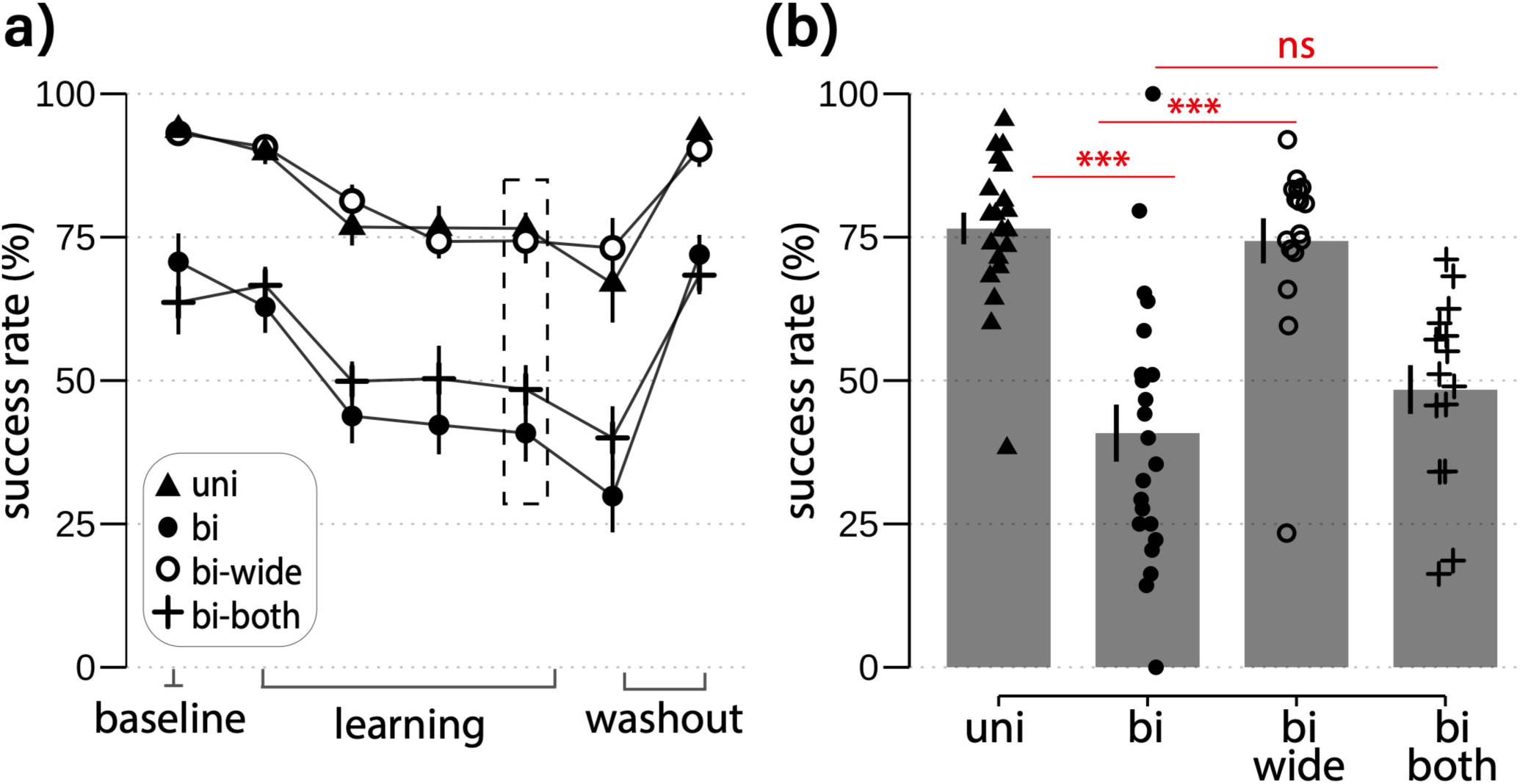
**(a)** Success rates by epoch across baseline, learning, and washout phases for the four groups. Unimanual group was more successful than bimanual group. **(b)** At the end of the learning phase (end plateau), success rates improved substantially when target was widened (bimanual-wide), but less so when both hands were perturbed (bimanual-both).

Widening the target substantially improved success rates in the bimanual-wide group, when compared to the bimanual group, across all experimental phases (Fig 2a, open circles) including at the end of the learning phase (Fig 2b, mean difference from the bimanual group = 37.6 ± 5.8%, z = 6.5, p<0.001), achieving similar success rates as the unimanual group. In contrast, removing interlimb conflict in the bimanual-both group did not improve success (Fig 2a, crosses), with success rates being similar to the bimanual group (Fig 2b, mean difference = 9.8 ± 6.6%, z = 1.5, p=0.13).

Thus, relaxing spatial precision demands, rather than eliminating interlimb conflict, restored success in the bimanual group to the level of the unimanual group.

### Tilt compensations were eased by removing interlimb conflict, not relaxing precision demands

Although plate movement was similar between the unimanual and bimanual groups during the learning phase (Fig 3a-b, uni: 29.0 ± 1.1 vs bi: 28.8 ± 1.1 cm, z = 0.45, p=0.65), hand movement patterns that led to this plate displacement diverged significantly between the uni- and bimanual groups as learning progressed.

**Figure 3.**
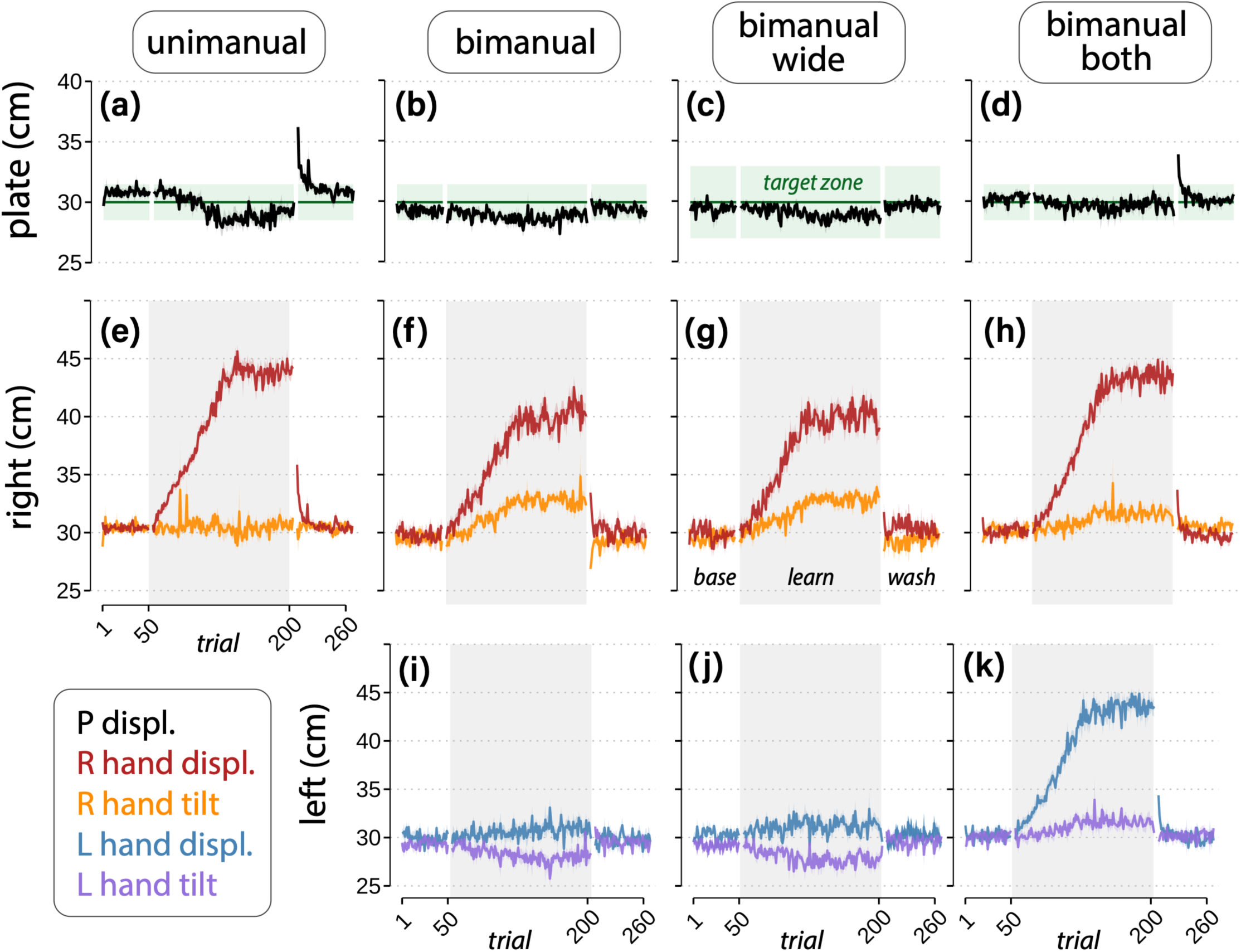
Trial-by-trial group averages for plate and hand displacement across the four groups. **(a–d)** Plate displacement (black) across all 260 trials for the unimanual (a), bimanual (b), bimanual-wide (c), and bimanual-both (d) groups. Green shading indicates the target zone (3 cm high in a, b, and d; 6 cm high in c). **(e–h)** Right-hand displacement (red) and right-hand tilt compensation (orange) across the four groups. **(i–k)** Left-hand displacement (blue) and left-hand tilt compensation (purple) across the four groups. Gray shading indicates the learning phase (ramp + plateau, trials 51–200); unshaded regions indicate baseline (trials 1–50) and washout (trials 201–260). The unimanual group performed the task with a single hand and thus has no corresponding left-hand panel. Tilt compensation traces (orange and purple) are offset by 30 cm for visualization purposes. Shading around each trace represents SEM; base = baseline, learn = learning phase (ramp + plateau), wash = washout.

Both groups gradually increased right-hand displacement in response to the visual perturbation (Fig 3e-f, red traces), but by the end of learning the unimanual group had raised the right hand significantly higher than the bimanual group (uni: 44.0 ± 2.1 vs bi: 40.1 ± 2 cm, z = 5.9, p<0.001) and did so without tilting the plate (orange trace). The bimanual group instead relied on compensatory movements, tilting the plate up with the right hand while increasing left-hand displacement (Fig 3j, blue trace) and tilting the left side down (purple trace) to keep the plate level. These compensations developed progressively over trials, with the bimanual group’s right-hand tilt compensation exceeding that of the unimanual group (uni: +0.45 ± 1.3 cm vs bi: +2.7 ± 1.3, z = 4.7, p<0.001).

Widening the target did not change this movement pattern. Despite the improved success rates reported above, plate (Fig 3c, 29.1 ± 0.84 cm) and right-hand displacement (Fig 3g, 40.1 ± 1 cm) by the end of learning were similar between the bimanual and bimanual-wide groups (plate: z = 0.77, p=0.44; right hand: z = 0.08, p=0.94). The compensatory right-hand tilt persisted and was similar to the bimanual group (+3 ± 1 cm, z = 1.5, p=0.13), as was the left-hand displacement (31.2 ± 1 cm, z = 0.58, p=0.56) and the opposing left-hand tilt (–2.2 ± 1.3 cm, z = 0.78, p=0.44).

Removing interlimb conflict, in contrast, altered the movement pattern. Plate displacement was slightly higher in the bimanual-both group (Fig 3d; 29.8 ± 0.76 cm) compared to the bimanual group by the end of learning (mean difference = 1 ± 0.34 cm; z = 2.9, p=0.003) but was achieved through significantly greater right-hand displacement (Fig 3i, 43.5 ± 1.06 cm, mean difference = 3.4 ± 0.6cm, z = 5.62, p<0.001) rather than compensatory tilt. Accordingly, the right-hand compensatory tilt was significantly reduced relative to the bimanual group (mean difference = 1.2 ± 0.4 cm, z = 3.15, p=0.002), though not eliminated when compared to the unimanual group (mean difference = 1.05 ± 0.4 cm, z = 2.6, p=0.008). As expected, left-hand displacement increased to match the right hand (Fig 3k, 43.6 ± 2.19 cm), while left-hand tilt reversed direction: rather than tilting downward as in the bimanual group (–2.1 ± 1.5 cm), it tilted upward in the bimanual-both group (+1.6 ± 1.5 cm, z = 7.1, p<0.001), moving in the same direction as the right-hand tilt instead of opposing it as it did in the bimanual group to level the plate.

Thus, while relaxing precision demands improved performance, it did not alter the underlying movement pattern, removing interlimb conflict allowed participants to rely more on right-hand displacement and less on tilt compensation, though the compensatory strategy was not fully eliminated.

### Removing interlimb conflict rescued plate aftereffects, but relaxing precision demands did not

During washout, the unimanual group showed an aftereffect in plate displacement that decayed slowly (Fig 4a, black trace, 33.1 ± 2.1 cm, z = 5.7, p<0.001), whereas the bimanual group showed no aftereffect (Fig 4b, black trace, 29.5 ± 2.1 cm, z = 0.24, p=0.81), consistent with our prediction that the bimanual group would adapt less (mean difference = 3.7 ± 0.56, z = 6.7, p<0.001).

**Figure 4.**
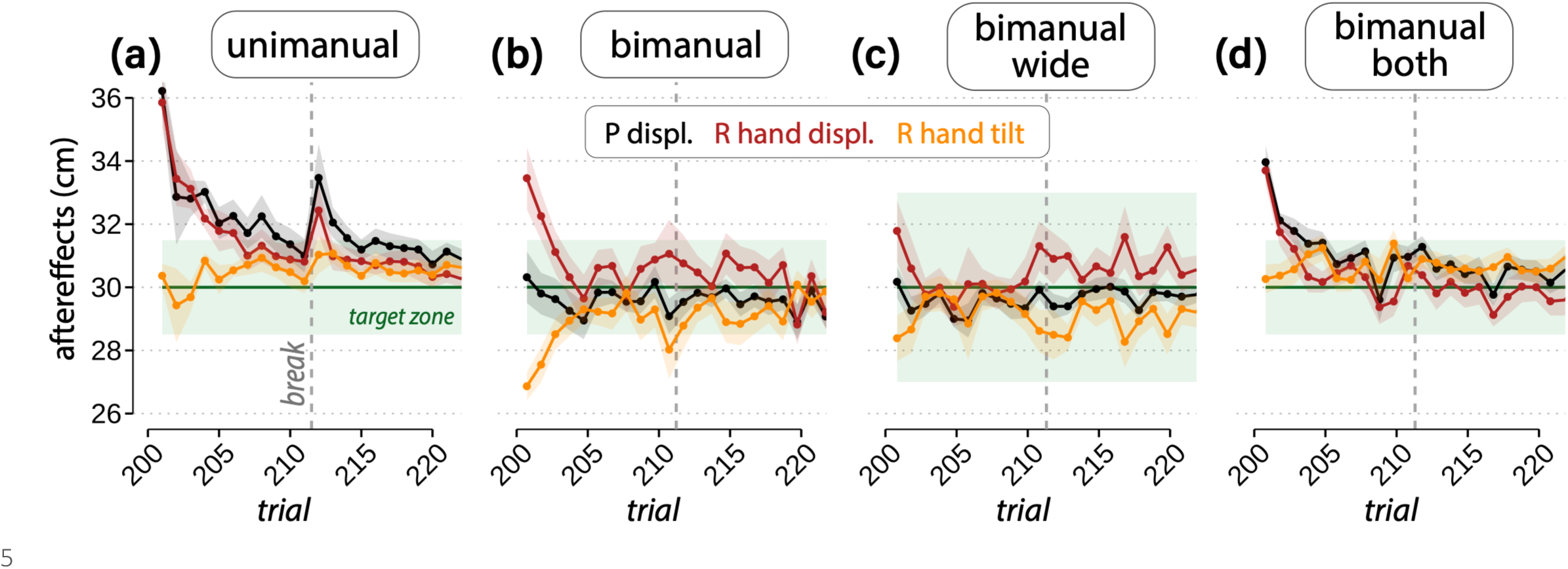
Aftereffects in plate and right-hand displacement and reactive tilt compensations in early washout across the four groups: **(a)** unimanual, **(b)** bimanual, **(c)** bimanual-wide, **(d)** bimanual-both. Trial-by-trial group averages for the first 20 trials of washout. Shading around each trace represents SEM; base = baseline, learn = learning phase (ramp + plateau), wash = washout.

A similar pattern held for the right hand. The unimanual group’s right-hand aftereffect (Fig 4a, red trace, 33.1 ± 2 cm, z = 4.6, p<0.001) washed out gradually and reappeared after a break (vertical dashed line), whereas the bimanual group’s right-hand aftereffect (Fig 4b, red trace, 31.3 ± 2.1 cm, z = 2.8, p=0.005) was smaller (mean difference = 3.9 ± 0.66 cm, z = 5.9, p<0.001) and more transient. The bimanual group’s right-hand tilt compensation countered the right-hand displacement (Fig 4b, orange trace, –1.8 ± 1.6 cm, z = 3.56, p<0.001), explaining the absence of a plate aftereffect. These tilt adjustments were present on the very first trial after the perturbation was removed, suggesting they were feedback-dependent rather than a retained adapted motor pattern. Of note, even in the unimanual group, where aftereffects were larger, they were still incomplete, i.e., only a portion of the learned right-hand displacement was expressed as aftereffects (44cm at the end of learning vs 33.1cm at the start of washout). This finding indicates that even in the unimanual group, participants were using a combination of feedback, strategy, and adaptation as opposed to adaptation alone.

Widening the target did not rescue either plate or right-hand aftereffect. Similar to the bimanual group, the bimanual-wide group showed no plate aftereffect (29.6 ± 0.97 cm) and similar magnitude right-hand aftereffects (Fig 4c, right-hand: 30.3 ± 1.9 cm; right tilt: 0.72 ± 1.9 cm). Aftereffects were of similar size across left-hand parameters (left displacement: 29.4 ± 1.5 cm; left tilt: 0.72 ± 1.9 cm; p>0.05 for both). Thus, relaxing precision demands was insufficient to restore adaptation, despite yielding higher success rates during learning.

Removing interlimb conflict, in contrast, rescued the plate aftereffect, but through a different mechanism than in the other groups. The bimanual-both group showed a significantly larger plate aftereffect than the bimanual group (Fig 4d, black trace, mean difference = 2.8 ± 0.46 cm, z = 6.1, p<0.001), though still smaller than the unimanual group (mean difference = 1.04 ± 0.45 cm, z = 2.3, p=0.02). This was not because the right-hand displacement aftereffect itself was larger; in fact, it did not differ significantly from the bimanual group (Fig 4d vs 4b, red trace). Instead, unlike the bimanual and bimanual-wide groups, where opposing right- and left-hand tilt cancelled out the residual plate displacement aftereffect (as described above), tilt in the bimanual-both group persisted in the same direction as displacement in both hands. As a result, the small hand aftereffect was not cancelled but preserved, and plate aftereffects were amplified.

Figure 5 summarizes these patterns across all three phases (baseline, end of learning, washout) for all four groups. The unimanual group showed the clearest signature of adaptation: substantial right-hand displacement with little-to-no tilt compensation during learning, and correspondingly large plate and right-hand aftereffects during washout (Fig 5a, e-f). The bimanual group showed the opposite pattern: smaller right-hand displacement, larger tilt compensation, and minimal aftereffects (Fig 5b, e-f); while its left hand changed comparatively little across phases (Fig 5i-j, blue and purple), consistent with the left hand serving a compensatory rather than an adaptive role.

**Figure 5.**
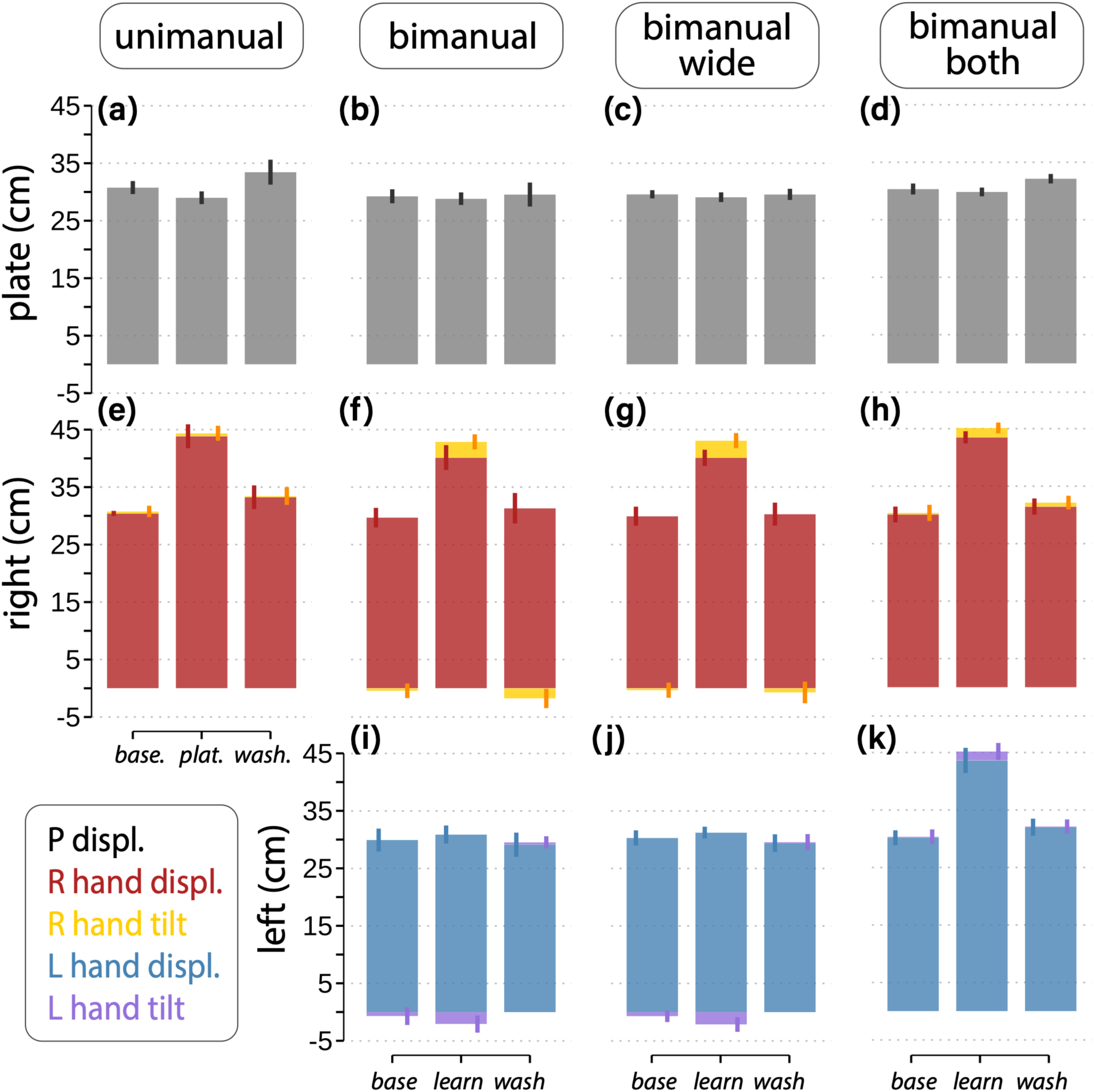
Stacked summary bar plots show mean ± SD plate displacement **(a–d)**, right-hand displacement (red) with tilt compensation (orange) stacked to show each component’s contribution to total movement **(e–h)**, and left-hand displacement with tilt compensation in the bimanual groups **(i–k)**, across baseline, end plateau, and early washout epochs. The **(a)** unimanual and **(d)** bimanual-both groups showed significant aftereffects in plate displacement in washout, whereas the **(b)** bimanual and **(c)** bimanual-wide groups did not. Right-hand tilt emerged by the end of plateau in the **(f)** bimanual group but not in **(e)** the unimanual group. **(g)** Tilt did not change when the target was widened but **(h)** reduced when both hands were perturbed. Left-hand tilt also emerged by the end of plateau **(i)** in the bimanual group but was in the opposite direction to that of the right-hand. Left-hand tilt was in the same direction as the right hand when **(k)** both hands were perturbed.

The two manipulations dissociated why the bimanual group adapted less. Widening the target (bimanual-wide) relaxed the precision demands that distinguished success between the unimanual and bimanual groups, and success rates recovered accordingly, but the underlying strategy did not change. Right-hand displacement, tilt compensation, and aftereffects all remained similar to the narrow-target bimanual group (Fig 5c, g). Applying the perturbation to both hands (bimanual-both) removed the interlimb conflict driving that compensation, and while success rates did not improve, the plate aftereffect was partially restored (Fig 5d), alongside a reduction in tilt compensation that was in the same rather than opposing direction for both hands (Fig 5h).

Together, these results suggest that the bimanual group’s reduced adaptation was driven less by the precision demands of the task and more by the interlimb conflict, and that resolving this conflict was sufficient to partially restore adaptation-like behavior reminiscent of the unimanual group.

### Aftereffects were associated with right-hand speed at the end of learning, not success or movement time

Having established these group-level differences, we next asked whether individual variation in performance and movement patterns predicted the magnitude of individual aftereffects. We examined whether success, movement time, or peak speed at the end of learning predicted aftereffects in plate and right-hand displacement during early washout. Across all groups, success at the end of learning did not correlate with either plate (Fig. 6a; r = 0.15, p = 0.21) or right-hand aftereffects (Fig. 6d; r = 0.092, p=0.44). Average movement time also did not correlate with plate (Fig. 6b; r = 0.16, p=0.19) or right-hand aftereffects (Fig. 6e; r = −0.23, p=0.055). However, right-hand peak speed at the end of learning was associated with larger plate (Fig. 6c; r = 0.27, p=0.022) and right-hand aftereffects (Fig. 6f; r = 0.44, p<0.001). Thus, participants who moved faster by the end of learning showed larger aftereffects.

**Figure 6.**
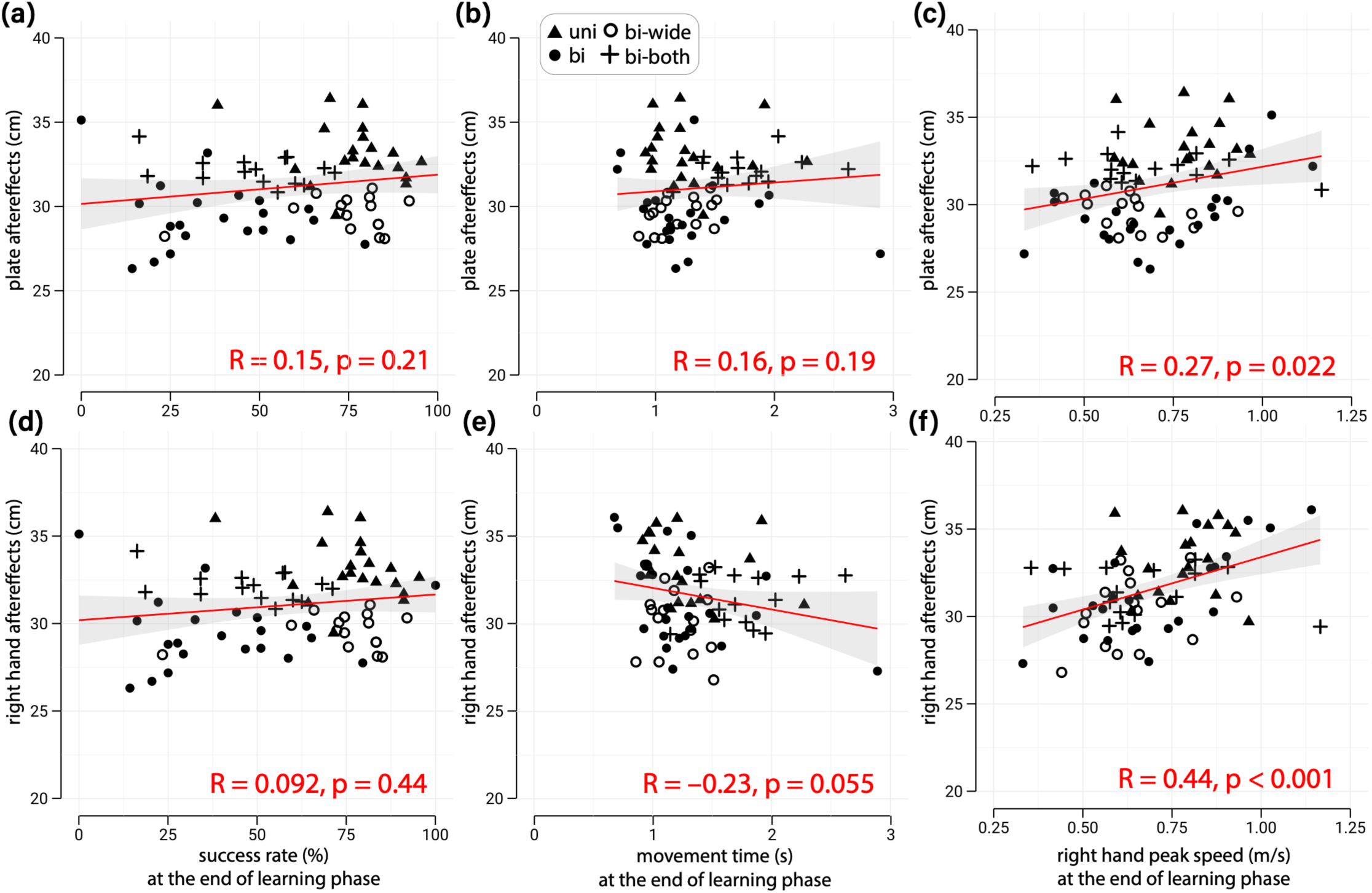
Although success rate and average movement time at the end of learning were not related to plate **(a, b)** or right-hand aftereffects **(d, e)**, faster right-hand peak speeds were associated with larger plate **(c)** and right-hand **(f)** aftereffects in early washout.

## DISCUSSION

We studied how task demands affect motor learning during unimanual and bimanual coordination. Given the unconstrained nature of the task (e.g., no speed requirements) and provision of full visual feedback, we anticipated that adaptation would be reduced compared to more constrained planar reaching tasks used in traditional motor learning studies. This was the case: aftereffects in the plate and right-hand movements accounted for only a portion of the displacement necessary to complete the task and varied in size across groups. Unimanual movements showed the greatest aftereffects whereas bimanual conditions showed little to no aftereffects depending on the task. Two factors seemed to influence this shift during bimanual tasks: precision demands and interlimb sensory conflict. Relaxing precision demands improved performance but did not reduce strategic compensation or bring back aftereffects. Reducing interlimb conflict, however, through bilateral perturbation, substantially cut compensation and partially restored aftereffects. These results show that interlimb conflict, not precision demands, shifts the balance between feedback control, adaptation, and other compensatory strategies in bimanual coordination. Notably, end-of-learning success did not correlate with greater adaptation, but movement speed did.

The differences between unimanual and bimanual groups are consistent with previous work showing that these contexts involve partially overlapping but distinct neural control processes.^23,24^ However, prior studies examined separate-goal bimanual actions, where each hand moved to different targets (amplitudes or directions) or at different times. Less is known about adaptation when both hands control a single shared object. Shared bimanual control introduces unique challenges: mapping task goals to commands for each hand when multiple hand configurations can achieve the same outcome and assigning responsibility for errors when there is interlimb conflict from unilateral perturbation. Participants in the bimanual group redistributed control between hands and across multiple joints, with tilt compensations that were in opposite directions between hands. The bimanual group also showed minimal aftereffects in plate displacement (Fig 4)— consistent with exploiting multiple motor solutions—and greater reliance on feedback corrections with smaller right-hand aftereffects, suggesting a greater shift from adaptation to feedback corrections compared to the unimanual group.

Previous work found that when both hands control a shared cursor and one hand experiences the perturbation, both hands adapt to the disturbance.^14,15^ Our findings differ: we observed no left-hand aftereffects (Fig 3i-j) except when directly perturbed as in the bimanual-both group (Fig3k). A likely explanation is clearer error attribution in our task: side-to-side plate tilt provided an alignment error that identified the right hand as the source of error, whereas single-point cursors convey only position error, leaving error attribution ambiguous and prompting both hands to adapt. This suggests that when alignment errors are available, feedback control can be distributed across hands while adaptation remains hand specific. It also raises the possibility that redundant motor solutions are exploited differently depending on the error signal available.^8^

To isolate how endpoint precision and interlimb conflict influenced bimanual adaptation, we conducted two control experiments. If precision requirements drive the shift away from adaptation, relaxing target requirements (wide target condition) should reduce feedback-driven corrections and strategic compensation, resulting in larger aftereffects. If interlimb conflict drives this shift, restoring sensory consistency (bilateral perturbation condition) should reduce the need for compensatory strategies and restore adaptation.

Relaxing precision requirements by widening the target improved success but was insufficient to eliminate compensatory strategies. Aftereffects in hand and plate displacement also remained unchanged despite improved success rates. Whether adaptation is sensitive to task success remains debated, with evidence both for^25–27^ and against^28–31^ such sensitivity, but here we did not find that the changes in success rate between the bimanual and bimanual-wide groups impacted adaptation as measured by the aftereffects. Our findings remained consistent even when we pooled data across all four groups: success rate at the end of learning did not correlate with plate or right-hand aftereffects (Fig 6a, d). Instead, right-hand peak speed at the end of learning correlated with aftereffects (Fig 6c, f). Recall the encyclopedia analogy: a box lifted quickly and smoothly, without correction, is a sign that the brain has learned to anticipate its weight in advance. In the same way, faster right-hand movements at the end of learning likely reflect the ability to better estimate the motor command needed to lift the plate and thus correlates more closely with aftereffects. Notably, despite reduced target errors in the bimanual-wide condition, compensatory hand adjustments persisted into washout. This pattern indicates these compensations may be driven by more than just target precision, and that precision demands alone cannot eliminate these strategies when sensory mismatch between hands remains.

When both hands experienced matched perturbations in the bimanual-both group, tilt compensations were substantially reduced and plate aftereffects increased to levels similar to the unimanual group. The interlimb coordination pattern also changed in the bimanual-both group compared to the bimanual group: in the bimanual-both group both right- and left-hand tilt compensations are in the same direction during learning whereas in the bimanual group right and left-hand tilt compensation are in the opposite direction. Similar flexibility has been observed in other bimanual tasks where the unperturbed hand moved either opposite to or in the same direction as the perturbed hand depending on task goals.^8^ Together, these findings demonstrate that removing interlimb conflict rescued plate aftereffects, whereas relaxing precision demands did not.

It should be noted that current VR technology introduces constraints that affect naturalism. VR systems introduce visual delay. Our estimated delay was 45 to 68 ms, consistent with other VR-based hand tracking studies.^32,33^ Although such small delays are thought to minimally impact presence in VR,^34,35^ we know that even small sub-100ms delays can impact adaptation.^36,37^ However, because this delay was similar across all experimental conditions, we can be more confident that learning differences likely arise from the experimentally manipulated task features rather than VR latencies. Current commercially available VR systems, like ours, also lack the haptic and kinetic feedback present in real-world object manipulation. For bimanual tasks, in particular, forces are not mechanically transmitted through the shared object in VR, potentially creating a decoupled environment. The absence of mechanical coupling and the greater visual attention typical of VR environments ^38,39,40^ may amplify reliance on visual feedback and tilt compensations. Recent advances in haptic-integrated VR systems may help address some of these limitations in the future.^41,42^

We chose a 3D vertical-axis lifting task because every day functional actions nearly always require accounting for gravity. While from a computational perspective, this may add complexity to the control problem, our goal here was to first characterize how such features of everyday bimanual tasks might constrain the adaptation process. Our control experiments (bimanual-wide and bimanual-both) partially addressed these challenges by systematically manipulating specific task features while holding other factors constant. Our interpretations regarding the balance between adaptation, feedback-driven corrections, and strategic compensation are based on empirical measurements (i.e., aftereffects, tilt) rather than model-derived estimates. Future studies could incorporate no-vision catch trials as an alternative approach to quantifying adaptation. Finally, we tested only neurotypical young adults in a single session, leaving open questions about retention, generalization, and clinical applicability.

In future work, we plan to test the feasibility of this VR-based bimanual training in stroke survivors. Our findings have implications for how we might design the task to focus on error-based learning or the better use of feedback or compensatory strategies for the impaired arm. For example, our results, particularly on restoring sensory consistency between the hands, suggest that error-based training of the impaired arm should account for existing asymmetries between the paretic and non-paretic arm in designing the perturbation to be applied to both hands. Key questions remain: Would stroke survivors, who often rely on compensatory strategies,^43^ similarly recruit additional degrees of freedom such as the trunk? Would such compensation interfere with or support adaptation of the affected arm? How well does learning generalize to real-world bimanual tasks? Gamified VR training has the potential to increase engagement and facilitate high-volume practice in clinic and at home.^44–47^ Whether error-based bimanual training in VR can improve function in clinical populations like stroke is a critical next question.

## METHODS

### Participants

Seventy-three neurotypical volunteers participated in one of four experimental groups (unimanual: n = 20; bimanual: n = 22; bimanual-wide: n = 16; bimanual-both: n = 15; see descriptions below). All participants were naïve to the paradigm and completed one of the four conditions. No formal power analysis was undertaken. Sample size was guided by pilot data and similar previous real-world motor learning studies.^20,48,49^ In accordance with the Declaration of Helsinki, study procedures were approved by the Johns Hopkins Medical Institutional Review Board (IRB00276939). All participants provided informed consent. Handedness was self-reported by participants. Sixty-nine of the 73 participants (94.5%) reported being right-handed.

### Setup and task

The task was implemented using the MovementVR platform^50^ on a Meta Quest 2 VR headset with real-time hand tracking via a LEAP Motion controller. Hand tracking sampled on average at 90 Hz (range: 75-120). Using high-speed video analysis at 240 fps the visual delay from physical hand movement onset to the PC monitor display was measured at 33.3 ± 4.2 ms. Accounting for Quest Link cable transmission, the total latency to the headset display was estimated at 45 to 68 ms. Leap Motion positional accuracy is <1 cm under optimal conditions.^51,52^ Data collection occurred in well-lit rooms to ensure optimal tracking quality.

Participants lifted a virtual plate of grapes with both hands transporting it to a target perch at eye level. The target perch was surrounded by an orange zone requiring precise plate placement. To account for individual differences in seated height, the setup was dynamically calibrated to each participant. Upon donning the headset, the initial position of the headset defined the center of the local tracking space (x = 0). The target perch was spawned at this initialized eye level, and the plate’s starting position was placed centered and 30 cm directly below the target.

Participants then lifted the plate to the target zone with full visual feedback of their hands and the plate. Trials ended when the plate entered the target zone, with binary audiovisual feedback indicating success (green flash and a chirp) or failure (red flash and a squawk). Total trial time was capped at 30 seconds. Instructions emphasized keeping the plate level and reaching the target accurately. If the plate tilted beyond ∼20°, grapes would slide off and the trial was unsuccessful. The virtual environment utilized Unity’s physics engine with a friction coefficient of 0.6 between the grapes, plate, and hands, allowing all objects to move freely in three dimensions; if the plate tilted beyond ∼20°, the grapes would slide off, resulting in an unsuccessful trial.

### Learning paradigm

The experiment consisted of 260 trials across three phases (Figure 1D). During baseline (50 trials), virtual hand movements matched real movements. In the learning phase (150 trials), visual gain was gradually reduced from 100% to 65% over 75 trials,^53^ then held constant for 75 trials. Gain perturbations were applied to the y-position of the wrist at plate contact. A washout phase (60 trials) restored normal visual feedback to examine aftereffects. Rest breaks occurred every 30 trials to minimize fatigue.

### Experimental groups

Four experimental conditions were tested to examine different factors affecting bimanual motor learning:

1. *Unimanual*: Participants used only their right-hand to lift a 13 cm diameter plate to a 3 cm target zone. Visual gain of the right-hand was reduced from 100% to 65% during adaptation.
2. *Bimanual*: Participants used both hands to lift a 26 cm diameter plate to a 3 cm target zone. Visual gain of the right-hand was reduced from 100% to 65% during adaptation.
3. *Bimanual-wide*: Identical to bimanual, but with a larger 6 cm target zone to reduce precision demands.
4. *Bimanual-both*: Identical to bimanual, but visual gain applied to both hands (100% to 65%) to eliminate interlimb conflict.

The four versions of the game (bimanual-narrow, unimanual-narrow, bimanual-wide, bilateral-perturbation) used in this study can be found here: https://osf.io/8wbkh/ as PC executable applications.

To make task constraints more comparable between the bimanual and unimanual group, we adjusted both the plate size and success criteria. The unimanual group used a smaller 13 cm plate (for more stable single-handed control) and needed only the plate center to reach the target zone. The bimanual group used a larger 26 cm plate but required both plate edges to reach the target zone. Success criteria also differed between unimanual and bimanual conditions. In unimanual, only the plate center needed to reach the target zone. In bimanual conditions, both left and right edges of the plate needed to reach the target zone to ensure participants experienced the full perturbation magnitude and maintained a level plate. This stricter criterion for the bimanual group ensured that both groups faced similar constraints on the range of acceptable plate alignment, thereby limiting redundancy in task solutions comparably across conditions.

### Data processing and kinematic measures

Primary analysis focused on the y-axis (lifting direction). Trials with plate displacement, less than 50% of the required distance were excluded, as were trials with technical errors (e.g., occasional loss in hand tracking, LEAP disconnection) or accidental plate drops (e.g., knocked over or flipped by hand on return). Time signals were linearly interpolated to yield 1000 samples. Hand and plate position data were filtered using a 4th-order Butterworth filter (5 Hz cutoff) and resampled to 1000 points via spline interpolation. Raw position data were corrected by subtracting initial values to normalize movement onset to zero. Velocity was calculated by differentiating position signals and smoothed for plotting using robust local regression (MATLAB Curve Fitting Toolbox: *smooth* function, ‘rloess’ method, 0.05 span). Movement onset was defined as first timepoint at which plate moved; offset as when the plate reached maximum height.

For visualization and statistical analysis, the following epochs were defined in advance: baseline (last 10 trials before perturbation), ramp phase (first and last 25 trials), plateau phase (first 25 and last 50 trials), and washout (first 5 and last 10 trials). Larger windows were used where behavior was stable but noisy; smaller windows where rapid changes were expected.^54^ Key measures included: (1) Success rates calculated as frequency of reward outcome within predefined epochs. (2) Plate, right- and left-hand displacement measured as the maximum vertical displacement from movement onset to offset. (3) Right- and left-hand compensation calculated as the difference between plate center displacement and seen right wrist displacement, capturing local adjustments by tilt or rotations. As noted earlier, our primary adaptation metric was aftereffects in plate and right-hand displacement in early washout.

### Statistical analysis

All statistical analyses were performed in R (version 4.5.1) and visualization using the *ggplot2* package (version 4.0.0). Statistical significance was set at α = 0.05 for all analyses. All measures were summarized as subject-wise means for predefined epochs for further analyses. Primary comparisons focused on end of plateau (end of adaptation) and start of washout (aftereffects). Missing trials were removed from analysis; at the end of adaptation, participants were missing a median of 4 trials (IQR: 2–7) out of 50; missingness at start of washout was negligible. All data reported in results and figures are as mean ± standard error unless otherwise specified.

A priori hypothesized group differences in epoch-averaged measures were assessed using robust linear models to minimize the influence of outliers (using the ‘rlm’ function in the *MASS* package version 7.3.65), which downweights outliers rather than excluding them.^55^ Comparisons included: unimanual versus bimanual for success rate, right-hand displacement, hand compensation, and aftereffects; bimanual versus bimanual-wide for success rate, peak speed, movement time, aftereffects, and hand compensation; and bimanual versus bimanual-both perturbation for hand compensation and plate aftereffects. To determine whether hand compensation was significantly different from zero in each group, estimated marginal means were tested against a null value of zero. An additional comparison examined plate aftereffects between the unimanual and bimanual-both perturbation groups to assess whether perturbing both hands restored adaptation toward unimanual levels. Estimated marginal means and pairwise contrasts (using the *emmeans* package version 1.11.2.8) ^56^ were computed, with Wald z-statistics reported based on asymptotic standard errors.

We used Spearman rank-order correlation coefficients to examine whether success rate, average movement time, and right-hand peak speed at the end of learning were associated with aftereffects in plate and right-hand displacement in early washout.

## Author Contributions

R.V. designed the study, collected and analyzed the data, performed statistical analyses, prepared all figures, and wrote the manuscript. C.R. developed the VR task methodology, contributed to study design and data interpretations. L.A.M. contributed to study design and data interpretation. A.J.B. secured funding, supervised the study, and contributed to study design and data interpretation. All authors reviewed and revised the manuscript.

## Data and Code Availability

Data and code to reproduce all the figures presented in this manuscript are available here: https://github.com/rinivarg/VRAdapt-Adults/tree/main

## Acknowledgements

This work was supported by the following funding sources: grants from the National Institutes of Health T32 HD007414 and R35 NS122266 to A.J.B. We thank Naser Al-Fawakhiri for helpful comments on earlier versions of this manuscript.

